# Controlling the flow of interruption in working memory

**DOI:** 10.1101/2020.09.08.288027

**Authors:** Nicole Hakim, Tobias Feldmann-Wüstefeld, Edward Awh, Edward K Vogel

**Affiliations:** Department of Psychology, University of Chicago, Chicago, IL; Institute for Mind and Biology, University of Chicago, Chicago, IL; Department of Psychology, University of Southampton, Southampton, UK; Grossman Institute for Neuroscience, Quantitative Biology, and Human Behavior, University of Chicago, Chicago, IL

**Keywords:** working memory gating, suppression, interruption, distraction

## Abstract

Visual working memory (WM) must maintain relevant information, despite the constant influx of both relevant and irrelevant information. Attentional control mechanisms help determine which of this new information gets access to our capacity-limited WM system. Previous work has treated attentional control as a monolithic process–either distractors capture attention or they are suppressed. Here, we provide evidence that attentional capture may instead be broken down into at least two distinct sub-component processes: 1) spatial capture, which refers to when spatial attention shifts towards the location of irrelevant stimuli, and 2) item-based capture, which refers to when item-based WM representations of irrelevant stimuli are formed. To dissociate these two sub-component processes of attentional capture, we utilized a series of EEG components that track WM maintenance (contralateral delay activity), suppression (distractor positivity), item individuation (N2pc), and spatial attention (lateralized alpha power). We show that relevant interrupters trigger both spatial and item-based capture, which means that they undermine WM maintenance more. Irrelevant interrupters, however, only trigger spatial capture from which ongoing WM representations can recover more easily. This fractionation of attentional capture into distinct sub-component processes provides a framework by which the fate of ongoing WM processes after interruption can be explained.

---

A critical feature of working memory is to protect internal representations from external interference. For example, when driving, the working memory representation of our route must be maintained despite irrelevant external interference, such as a flashing colorful billboard. Nevertheless, external information is sometimes relevant, and working memory must integrate this new information with our ongoing working memory representations. For example, a flashing sign warning us of a car accident ahead may capture our attention – but to our advantage. This new information allows us to update our working memory representation of our route in order to avoid traffic caused by the car accident. Attentional control mechanisms help determine which information gets access to our capacity-limited working memory system.

Models of attentional control suggest that attentional selection is based on a competitive process (Bundesen, 1990; Duncan & Humphreys, 1989) in which both goal-driven and stimulus-driven factors determine which information is selected from a given display (Awh et al., 2012; Corbetta & Shulman, 2002; Desimone & Duncan, 1995; Egeth & Yantis, 1997; Folk, Remington & Johnston, 1992; Itti & Koch, 2000; Jonies, 1981; Kastner & Ungerleider, 2000; Posner, 1980; Posner & Petersen, 1990; Wolfe, 1004). But can goal-driven attentional selection override stimulus-driven capture? Some previous research has suggested that salient interrupting information captures attention before it can be suppressed (Hickey et al., 2009; Liesefeld et al., 2017; Sawaki et al., 2012; Feldmann-Wüstefeld & Schubö, 2013). Based on this, they argue that attentional capture is obligatory because salient information needs to be processed before being discarded. Other research suggests that distractor suppression can prevent attentional capture (Feldmann-Wüstefeld et al., 2020; Gaspar & McDonald, 2014; Gaspelin et al., 2015, 2016; Gaspelin & Luck, 2018). This work has found that, for example, participants make fewer erroneous eye movements towards (Gaspelin et al., 2016) and are less likely to report the identity of (Gaspelin et al., 2015) a successfully suppressed salient distractor than a less salient distractor that was not suppressed. Although these appear to be conflicting perspectives, we suggest here that they might be reconciled by distinguishing between two distinct forms of attentional capture.

Attentional capture and suppression are often treated as a monolithic process: either a distractor captures attention or it is suppressed. However, recent work suggests that attention may include two distinct sub-component processes: attention to regions in space and representations of objects that occupy the attended regions (Hakim, Adam, et al., 2019). Therefore, we propose that involuntary attentional capture may also be broken down into at least two distinct sub-component processes: 1) Spatial capture, which refers to when spatial attention shifts towards the location of irrelevant stimuli 2) Item-based capture, which refers to when item-based representations of irrelevant stimuli are formed in working memory. Although spatial and item-based capture may have distinct operating principles, prior work has typically been unable to distinguish between these two forms of capture.

Thus, in the present work, we separately measured spatial and item-based capture using EEG activity. We used lateralized alpha power (8-12 Hz) to track spatial capture (Foster et al., 2015; Hakim, Adam, et al., 2019). This oscillatory signal has been shown to track attended hemifield (Hakim, Adam, et al., 2019) and has been shown to contain precise spatial information about attended stimuli (Foster et al., 2015, p. 201, 2017). To track item-based capture, we used the contralateral delay activity (CDA) and the distractor positivity (Pd). The CDA tracks the number of items maintained in working memory (Balaban & Luria, 2017; Luria et al., 2016), whereas the Pd tracks the suppression of irrelevant information (Burra & Kerzel, 2014; Feldmann-Wüstefeld & Schubö, 2013; Hickey et al., 2009), while also being sensitive to the number of irrelevant items that are presented (Feldmann-Wüstefeld & Vogel, 2019).

We used these EEG signals to assess how salient items with a sudden onset (interrupters) are processed when subjects are maintaining relevant information in working memory. In addition, we manipulated the task-relevance of the interrupters to compare how each type of attentional capture is influenced by goal-driven selection. Finally, while past work has typically focused on competition between simultaneously presented targets and distractors, here we focused on how interruptions influence the maintenance of items that have already been stably encoded into working memory. This provided the opportunity to obtain clear evidence regarding the degree to which interrupters elicited spatial and item-based attentional capture, and the distinct impact of task relevance on each form of capture.

Prominent models of attentional control assert that visually selected stimuli should automatically gain access to working memory at least for a short period of time (e.g., Bundesen et al., 2005). Moreover, previous research has shown that a sudden onset of salient but irrelevant information captures attention (Feldmann-Wüstefeld et al., 2015; Theeuwes, 2010) and that this negatively impacts ongoing working memory representations (Bisley & Goldberg, 2010; Hakim, Feldmann-Wüstefeld, et al., 2019; van Moorselaar et al., 2017). Does this disruption of working memory maintenance reflect an obligatory encoding of distracting information into working memory? To anticipate the results, we observed clearly distinct effects of task relevance on spatial and item-based attentional capture. Continuous tracking of alpha laterality showed that spatial attention was captured by interrupting stimuli, regardless of whether they were relevant or not. In sharp contrast, item-based attentional capture was completely determined by task relevance. Relevant interrupters were encoded into working memory, as shown by N2pc and CDA signals that tracked item individuation and working memory maintenance, respectively. By contrast, when the interrupters were irrelevant, a Pd was observed contralateral to the interrupters and no CDA was observed. Thus, irrelevant interrupters were not actively encoded into working memory, even though they clearly captured spatial attention. Thus, our findings offer a potential reconciliation of prior conflicting findings by showing that the encoding of interrupting information into working memory can be suppressed even when there is clear evidence that spatial attention has been captured.

## Materials & methods

### Experiment 1

In Experiment 1, we sought to determine how task-relevant versus task-irrelevant interrupters are processed. To this end, we presented memory array items along the midline and presented targets laterally. This allowed us to isolate the neural representations of the interrupters themselves. With this design, any lateralized signal, such as CDA or lateralized alpha power, should reflect the processing of the interrupters and not the memory array.

Previous research has shown that active representations may be required for the identification of relevant stimulus features (Mazza et al., 2007; Mcdonald et al., 2013). Therefore, we predicted that when participants had to discriminate the interrupters, they would be more likely to encode them into working memory than when the interrupters could be ignored. Accordingly, there should be a CDA following interruption when they discriminate the interrupters. Conversely, when participants could ignore the interrupters, we predicted that they would actively suppress them, as their features do not need to be identified. The distractor positivity (PD) has been shown to track suppression of irrelevant information (Burra & Kerzel, 2014; Feldmann-Wüstefeld & Schubö, 2013; Risa Sawaki & Luck, 2012). Therefore, we should expect to find a robust PD when participants ignore the interrupters, but not when they discriminate them.

#### Participants

Thirty novel volunteers, naïve to the objective of the experiment participated for payment ($15 USD per hour). Data from one participant was excluded from the analysis because of technical issue with the behavioral data file. Data from nine participants were excluded from the analysis because of too many artifacts that resulted in fewer than 150 trials in any condition. The remaining 20 participants (6 male) were between the ages of 21-31 (M = 23.5, SD = 3.3). Participants in all experiments reported normal or corrected-to-normal visual acuity as well as normal color vision. All experiments were conducted with the written understanding and consent of each participant. The University of Chicago Institutional Review Board approved experimental procedures.

#### Stimuli

All stimuli were presented on a gray background (~33.3 cd/m^2^). Cue displays showed a central fixation dot (0.2° × 0.2°). Memory displays showed four colored squares (1.1° by 1.1°, mean luminance 43.1 cd/m^2^) along the midline with a randomly jittered horizontal offset of maximally 0.55° (half of an object). Colors for the squares were selected randomly from a set of 11 possible colors (Red = 255, 0, 0; Green = 0, 255, 0; Blue = 0, 0, 255; Yellow = 255, 255, 0; Magenta = 255, 0, 255; Cyan = 0, 255, 255; Purple = 102, 0, 102; Brown = 102, 51, 0; Orange = 255, 128, 0; White = 255, 255, 255; Black = 0, 0, 0). No color was repeated. On 50% of trials, the retention interval display remained blank with a central fixation dot (0.2° × 0.2°). However, on the other 50% of trials, during the delay interrupters appeared laterally. On one side of the screen, four colored circles (25% of all trials) or four squares (25% of all trials) appeared during the delay. Items from these colors were chosen from the say 11 possible colors, but were never the same as the memory array items on a given trial. Additionally, these items had the same area as the items from the memory array. On the other side of the screen, four gray diamonds appeared (RGB= 80, 80, 80) at the same time as the colored circles/squares. These diamonds were the same area as the circles and squares and were luminance matched on average to the colored interrupters. We presented these gray diamonds so as to match the bottom-up visual stimulation on both sides of the screen. The hemifield in which the diamonds and the hemifield in which the colored circles/squares were presented were randomly selected each trial. All stimuli had the same area. Probe displays showed one colored square along the midline in the same location as one of the memory array items, randomly picked, in the original array. In 50% of the trials, the color of the square in the attended hemifield was identical (no change trial) to the memory display. In the remaining 50% of trials, it was one of the colors not used in the memory or interruption display (change trials).

#### Apparatus

Participants were seated with a chin-rest in a comfortable chair in a dimly lit, electrically shielded and sound attenuated chamber. Participants responded with button presses on a standard keyboard that was placed in front of them. Stimuli were presented on an LCD computer screen (BenQ XL2430T; 120 Hz refresh rate; 61 cm screen size in diameter; 1920 × 1080 pixels) placed at 74 cm distance from participants. An IBM-compatible computer (Dell Optiplex 9020) controlled stimulus presentation and response collection.

#### Procedure

Each trial began a memory display consisting of four colored squares along the midline appeared for 150 ms. Participants were instructed to memorize as many colored squares in the memory display as possible. Participants had to remember the items over a blank retention interval that contained a central fixation dot. The retention interval lasted 2,000 ms, regardless of whether an interruption appeared. In 50% of the trials, an interruption display appeared 500 ms after memory display offset for 150 ms. The interruption display consisted of four circles (50% of interrupted trials) or four squares (50% of interrupted trials) that appeared laterally. This display was visually balanced by four gray diamonds that appeared in the opposite hemifield. In half of the trials, participants were instructed to ignore interruption displays (Ignore block). In the other half of the trials, participants were instructed to discriminate the shape of the interrupting items (Discriminate block). After the retention interval, a probe display appeared until response. In both Ignore and Discriminate blocks, participants had to indicate whether the object at the probed location of the attended hemifield changed color (“?/” key) or did not change color (“z” key). In the Discriminate block, participants additionally performed a go-no-go task. Before responding to the probe, they had to press “space” to indicate that the interrupting objects were circles. If the interrupting objects were squares, they did not have to press any key. After participants responded, the next trial started after a blank inter-trial interval of 750 ms. Participants completed a total of 1600 trials (20 blocks of 80 trials), i.e. 800 trials with interruption and 800 trials without interruption. The first half of the experiment was always the Ignore blocks, and the second half of the experiment was always the Discriminate blocks. Information about average performance and a minimum break of 30 seconds was provided after each block. See Figure 1 for a visual depiction of the task.

**Figure 1.**
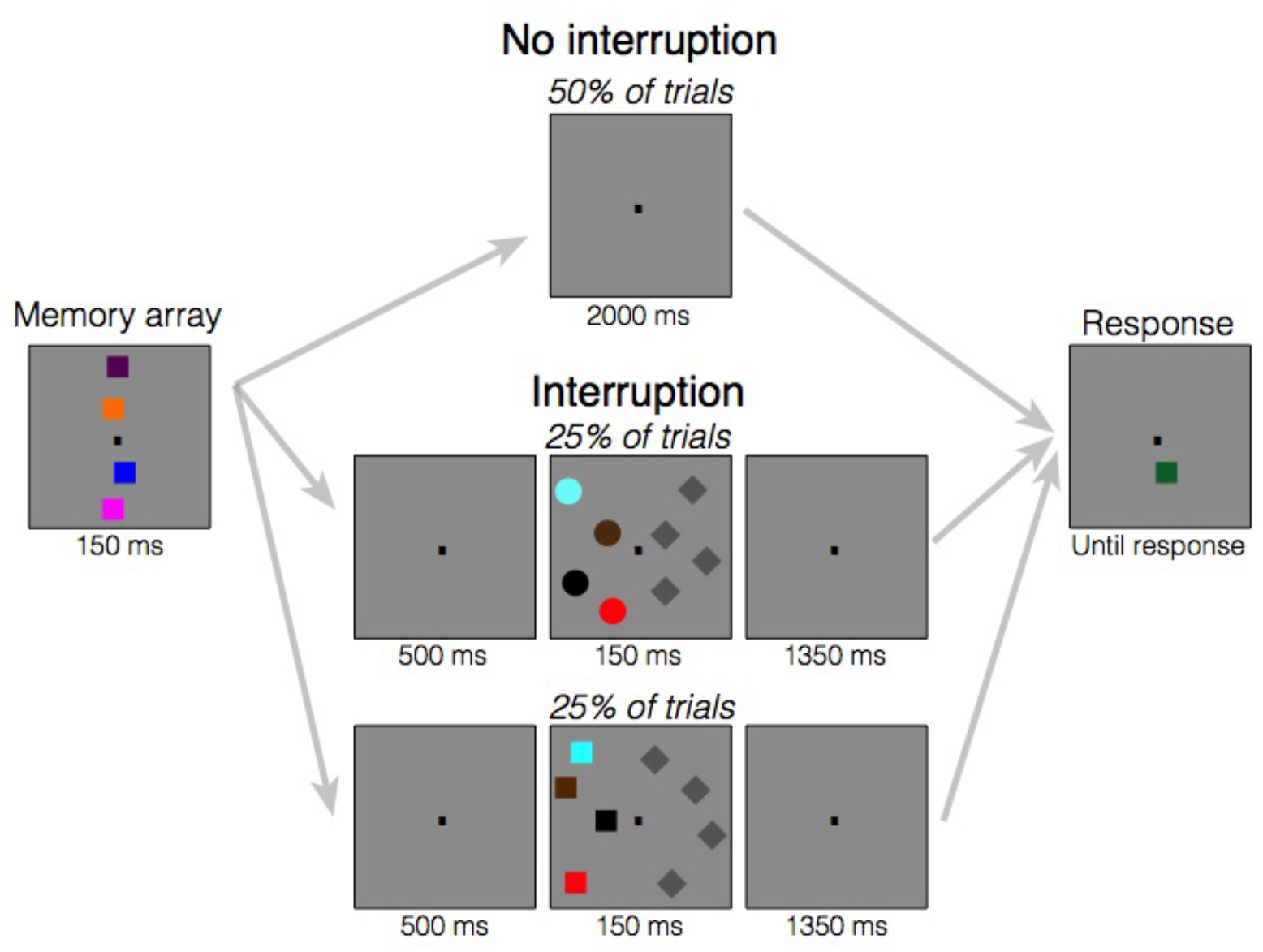
Task design for Experiment 1. At the start of each trial, the memory array appeared, which consisted of four colored squares along the midline. Participants were told to remember the colors and locations of these squares over the brief delay. Following the memory array, the screen went blank. Then, either the screen remained blank the entire delay (no interruption conditions) or the screen remained blank for 500 ms (interruption condition) followed by a series of interrupters presented laterally. These interrupters consisted of either four colored squares or circles on one side of the screen and iso-luminant gray diamonds on the other side of the screen. When participants were in the “Ignore” condition, they were told to always ignore these interrupting objects. When they were in the “Discriminate” condition, they were told to determine the shape of the colored stimuli (squares vs. circles) in order to report whether the stimuli were circles. They were told to withhold their response until the response screen appeared. Following interrupters, the screen then went blank for the rest of the delay. On the final screen, one square on the midline re-appeared and could either be the same color as the original square or it could be a different color. In both conditions, participants had to report whether the square on the cued side of the screen changed colors. In the “Discriminate” condition, participants additionally had to report whether the interrupters were circles, if there were interrupters on that trial.

We presented the interrupters in locations that did not overlap with the locations of the memory items to avoid visual masking. Importantly, the relative position of interrupters and targets matter in this kind of change detection tasks. When interrupters are presented laterally with targets on the vertical midline, the neural signature of sustained interrupter suppression can be isolated (CDAp). Conversely, when interrupters are presented on the vertical midline and targets are presented laterally, the neural signature of target processing can be isolated (Feldmann-Wüstefeld & Vogel, 2018). Accordingly, in Experiment 1, we were interested in the neural representations of the interrupters, so we placed the interrupters laterally. Thus, lateralized signals, such as CDA and lateralized alpha power, could be used to assess item-based and spatial capture elicited by the lateralized interrupters.

### Artifact rejection

We recorded EEG activity from 30 active Ag/AgCl electrodes (Brain Products actiCHamp, Munich, Germany) mounted in an elastic cap positioned according to the International 10-20 system [Fp1, Fp2, F7, F8, F3, F4, Fz, FC5, FC6, FC1, FC2, C3, C4, Cz, CP5, CP6, CP1, CP2, P7, P8, P3, P4, Pz, PO7, PO8, PO3, PO4, O1, O2, Oz]. FPz served as the ground electrode and all electrodes were referenced online to TP10 and re-referenced off-line to the average of all electrodes. Incoming data were filtered [low cut-off = .01 Hz, high cut-off = 80 Hz, slope from low-to high-cutoff = 12 dB/octave] and recorded with a 500 Hz sampling rate. Impedances were kept below 10kΩ. To identify trials that were contaminated with eye movements and blinks, we used electrooculogram (EOG) activity and eye tracking. We collected EOG data with 5 passive Ag/AgCl electrodes (2 vertical EOG electrodes placed above and below the right eye, 2 horizontal EOG electrodes placed ~1 cm from the outer canthi, and 1 ground electrode placed on the left cheek). We collected eye-tracking data using a desk-mounted EyeLink 1000 Plus eye-tracking camera (SR Research Ltd., Ontario, Canada) sampling at 1,000 Hz. Usable eye-tracking data were acquired for 20 out of 22 participants in Experiment 1 and 29 out of 30 participants in Experiment 2.

EEG was segmented offline with 2000 ms segments time-locked to memory display onset, including a 200 ms pre-stimulus baseline period. Eye movements, blinks, blocking, drift, and muscle artifacts were first detected by applying automatic detection criteria to each segment. After automatic detection (see below), trials were manually inspected to confirm that detection thresholds were working as expected. Participants were excluded if they had less than 150 correct trials remaining in any of the conditions. For the participants used in analyses, we rejected on average 12% of trials in Experiment 1, and 27% of trials in Experiment 2.

#### Eye movements

We used a sliding window step-function to check for eye movements in the HEOG and the eye-tracking gaze coordinates. For HEOG rejection, we used a split-half sliding window approach. We slid a 100 ms time window in steps of 10 ms from the beginning to the end of the trial. If the change in voltage from the first half to the second half of the window was greater than 20 μV, it was marked as an eye movement and rejected. For eye-tracking rejection, we applied a sliding window analysis to the x-gaze coordinates and y-gaze coordinates (window size = 100 ms, step size = 10 ms, threshold = 0.5° of visual angle).

#### Blinks

We used a sliding window step function to check for blinks in the VEOG (window size = 80 ms, step size = 10 ms, threshold = 30 μV). We checked the eye-tracking data for trial segments with missing data-points (no position data is recorded when the eye is closed).

#### Drift, muscle artifacts, and blocking

We checked for drift (e.g. skin potentials) by comparing the absolute change in voltage from the first quarter of the trial to the last quarter of the trial. If the change in voltage exceeded 100 μV, the trial was rejected for drift. In addition to slow drift, we checked for sudden step-like changes in voltage with a sliding window (window size = 100 ms, step size = 10 ms, threshold = 100 μV). We excluded trials for muscle artifacts if any electrode had peak-to-peak amplitude greater than 200 μV within a 15 ms time window. We excluded trials for blocking if any electrode had at least 30 time-points in any given 200-ms time window that were within 1V of each other.

### Behavioral data analysis

We separately analyzed performance for four separate conditions: trials without interrupters in the ignore block, trials with interrupters in the ignore block, trials without interrupters in the discriminate block, and trials with interrupters in the discriminate block. Performance was converted to a capacity score, K, calculated as N x (H-FA), where N is the setsize, H is the hit rate, and FA is the false alarm rate (Cowan, 2001). To compare performance between conditions, we used a two-way ANOVA with the within-subjects factors Interruption (interruption versus no interruption) and Relevance (ignore versus discriminate). All analyses were done with circle and square interrupters collapsed, since the circles and squares were equiprobable (circles: 50% of interrupted trials, squares: 50% of interrupted trials). We additionally ran twotailed follow-up t-tests when it was justified.

### Lateralized ERP analyses

Segmented EEG data was baselined from 200 ms to 0 ms before the onset of the memory displays. Artifact-free EEG segments were averaged separately for the two conditions when interrupters appeared (relevant versus irrelevant interrupters). Data was not analyzed for trials without interrupters because “laterality” was undefined in this condition. The difference between contralateral and ipsilateral activity for the electrode pair PO7/PO8 was calculated (i.e., the CDA), resulting in two average waveforms for each participant (one per analyzed condition). The average CDA amplitude was calculated for three time windows: before interruption onset (400-650 ms), and two windows following interruption offset (850-950 ms and 1050-1300ms). Previous research has shown that CDA amplitude should stabilize approximately 400 ms after memory array onset. To measure CDA after it stabilized, we chose a time window starting 400 ms after onset of the memory array. We wanted to measure an analogous time window following the onset of the interruption (i.e. 400 ms after interruption onset), which is why we chose the time window 1050-1300ms. For all of these time windows, we then compared the CDA across conditions with a paired-samples t-test. To measure the robustness of the CDA for each condition (reliable difference between contra- and ipsilateral activity), we also ran t-tests (against zero) for each time window and condition. These t-tests are two-tailed, unless otherwise stated.

In trials with interruption, we additionally analyzed the Distractor Positivity (Pd) and the N2pc. We used a data-driven approach (Feldmann-Wüstefeld & Vogel, 2018) to calculate the lateralized waveform (contra-minus ipsilateral to interrupters) for electrodes PO7/PO8, across participants and conditions. We determined the peak of the Pd and N2pc as the most positive or negative peak, respectively, 200 to 350 ms after interruption onset across both conditions. The average amplitude from 20 ms before to 20 ms after that peak was used for statistical analyses on the Pd and the average amplitude from 50 ms before to 50 ms after that peak was used for statistical analyses on the N2pc.

### Lateralized alpha power analysis

For the alpha power analyses, we did not baseline the segments. The raw EEG signal was band-pass filtered in the alpha band (8-12 Hz) using a two-way least-squares finite-impulse-response filter (“eegfilt.m” from EEGLAB Toolbox). Instantaneous power was then extracted by applying a Hilbert transform (‘hilbert.m’) to the filtered data. The resulting data were averaged separately for the two conditions when interrupters appeared (relevant versus irrelevant interrupters) and each laterality (contra-versus ipsi-lateral to cued hemifield) for the electrode pair PO7/PO8. Average alpha power was calculated for two of the same time windows as the CDA analysis: before interruption onset (400-650 ms), and post-interruption offset I (850-950 ms). Previous research has not investigated lateralized alpha power while participants maintained working memory representations that were presented centrally. Therefore, in this experiment, we did not have strong a-priori predictions about the timing of alpha power lateralization following the onset of lateralized interruptions. Additionally, even when relevant memoranda are presented laterally, alpha power takes up to 1,000 ms after memory array onset to become fully lateralized (Hakim, Adam, et al., 2019; Hakim, Feldmann-Wüstefeld, et al., 2019). Therefore, third time window that we analyzed extended to included up to 1,000 ms after interruption onset (1050-1650 ms). We then compared alpha power lateralization for each time window with a paired-samples t-test. To measure the robustness of alpha power lateralization for each condition (reliable difference between contra- and ipsilateral activity), we also ran t-tests (against zero) for each time window and condition. These t-tests are two-tailed, unless otherwise stated.

## Results

### Behavior

Participants remembered fewer items when they were interrupted (M=1.95, sd=0.61) than when they were not interrupted (M=2.66, sd=0.60). This difference was larger when the interrupters had to be discriminated (ΔM=1.001, sd=0.297) than when they could be ignored (ΔM=0.419, sd=0.293). This was evident from the significant interaction of Interruption and Discrimination (F(1,19)=55.725, p<0.001, η_p_^2^ =0.746), and from the significant follow-up paired-samples t-test (t(19)=-7.465, p<0.001). This t-test compared the difference between trials with and without interrupters in the Ignore and the Discriminate conditions. The main effects of Interruption and Discrimination were also significant (both p<0.001). Behavioral results for Experiment 1 are depicted in Figure 2.

**Figure 2.**
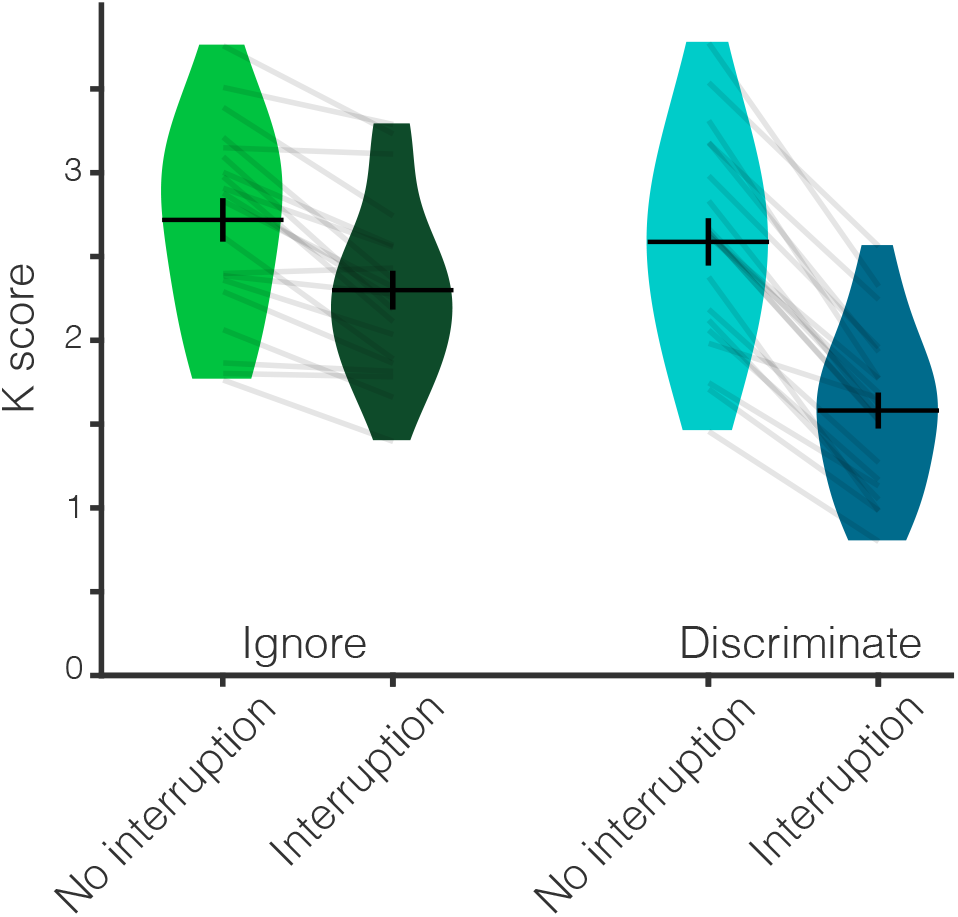
Behavioral results from Experiment 1. Behavioral performance (K score) across the four conditions. Participants remembered fewer items when they were interrupted than when they were not interrupted. This impact of interruption was larger when participants had to discriminate the interrupters than when they could ignore them. Average K score is represented by the horizontal black line and the black error bars reflect the standard error of the mean. The distribution of K scores in each condition for all participants is represented by the violin plots. Light gray lines connect data from one participant across conditions.

### Lateralized ERP

In this experiment, we analyzed lateralized alpha power, the contralateral delay activity (CDA), distractor positivity (Pd), and N2pc. For all of the below analyses, we calculated the difference between contralateral and ipsilateral activity (see Figure 3 for all neural results from Experiment 1) and then took the difference (contralateral – ipsilateral). We then analyzed this difference value for the two conditions when interrupters were presented (relevant versus irrelevant) to determine whether there were any lateralized differences between conditions at each time window. To determine whether the signals were significantly lateralized, we additionally calculated one-way t-tests for each condition.

**Figure 3.**
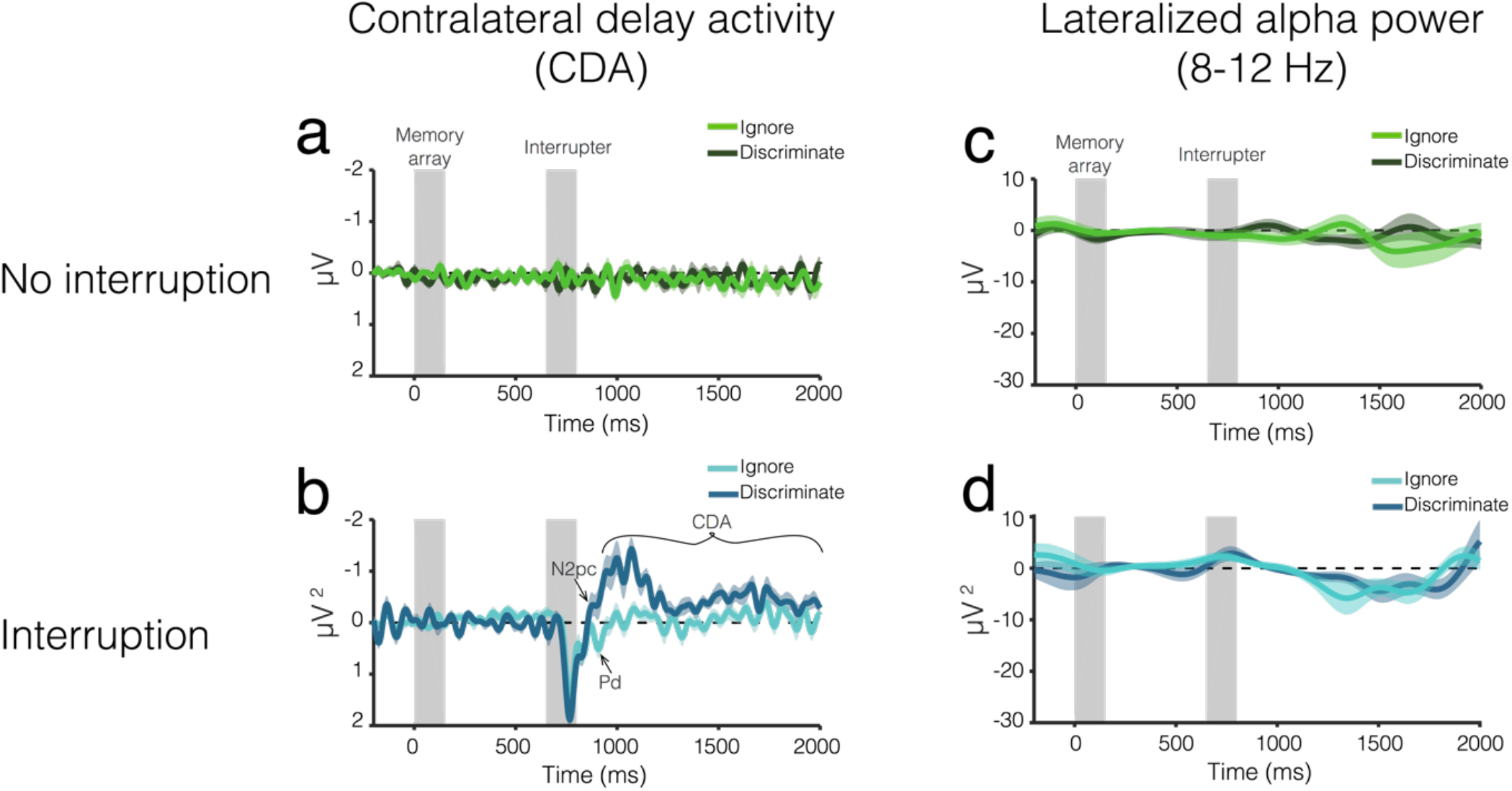
EEG results from Experiment 1. Average CDA amplitude over time for trials (b) with and (a) without interruption. The light color envelopes around each line represent standard error of the mean for each condition. The first vertical gray bar (0-150 ms) represents when the memory array was on the screen, and the second gray bar (650-800 ms) represents when the interrupters were on the screen, if there were interrupters on that trial. Lateralized alpha power over time for trials (d) with and (c) without interruption.

#### Pre-interruption (400-650 ms)

Before interruption onset (400-650 ms), the were no lateralized ERPs in either condition (Ignore: t(19)=-1.78, p=0.09, d=-0.40; Discriminate: t(19)=0.51, p=-0.62, d=0.11), and there was no difference between trials with relevant versus irrelevant interrupters (t(19)=-1.53, p=0.14, d=0.24). This is what we expected because the memory array was presented centrally and the lateralized interrupters had not yet appeared.

#### Post-interruption I (850-950 ms)

During this time window (equivalent to 200-300 ms post interruption, typically used for attention components), we observed a robust difference in lateralization between trials with relevant and irrelevant interrupters (t(19)=3.19, p=0.005, d=0.71). On trials with interrupters, the lateralized ERP was positive in the ignore condition (M=0.24±0.45) and negative in the discriminate condition (M=-0.44±0.98), suggesting the presence of an N2pc and Pd, respectively. To confirm this, we calculated the Pd and N2pc in specific a-priori time windows (see Methods for detail). One-way, one-sample t-tests against zero confirmed that there was a reliable Pd when interrupters were present and ignored (t(19)=2.34, p=0.02, d=0.53) and no Pd when interrupters were present and discriminated (t(19)=-1.65, p=0.94, d=-0.37). Additionally, there was a reliable N2pc when interrupters were present and discriminated (t(19)=-2.45, p=0.01, d=-0.55), but not when interrupters were presented and ignored (t(19)=2.37, p=0.99, d=0.53)

#### Post-interruption II (1050-1300 ms)

In this time window, there was a significant difference in lateralization between trials with relevant and irrelevant interrupters (t(19)=4.32, p<0.001, d=0.97). The CDA was more lateralized when the interrupters had to be discriminated (M=-0.59±0.55) than when interrupters needed to be ignored (M=-0.08±0.30). In fact, the CDA was reliable on trials with interrupters that needed to be discriminated (one-sample: t(19)=-4.81, p<0.001, d=-1.08), but not on trials with interrupters that could be ignored (one-sample: t(19)=-1.13, p=0.27, d=-0.25).

### Lateralized alpha power

#### Pre-interruption (400-650 ms)

Before interruption onset (400-650 ms), alpha power was not significantly lateralized in either condition (Ignore: t(19)=1.13, p=0.27, d=0.25; Discriminate: t(19)=-0.57, p=0.57 d=-0.13), and there was no difference between trials with relevant versus irrelevant interrupters (t(19)=0.991, p=0.33, d=0.22). This was expected because the memory array was presented centrally and the lateralized interrupters had not yet appeared.

#### Post-interruption I (850-950 ms)

Immediately following interruption (850-950 ms), alpha power was not significantly lateralized in either condition (Ignore: t(19)=1.20, p=0.23, d=0.27; Discriminate: t(19)=1.50, p=0.16, d=0.33), and lateralization did not vary between conditions (t(19)=-0.23, p=0.82, d=-0.05).

#### Post-interruption II (1050-1300 ms)

Towards the end of the trial (1050-1300 ms), alpha power was significantly lateralized in both conditions, consistent with a shift of spatial attention towards the interrupters (Ignore: t(19)=-2.33, p=0.031, d=-0.52; Discriminate: t(19)=-2.24, p=0.037, d=-0.50). Interestingly, there was no difference in lateralization between the two conditions (paired-samples t-test: t(19)=-0.89, p=0.40, d=-0.20).

### Conclusions

In Experiment 1, participants performed a WM change detection task with interrupters that appeared during the delay on a subset of trials. Behaviorally, participants remembered fewer items when they were interrupted than when they were not interrupted. This negative impact of interruption on behavior was larger when participants discriminated the interrupters than when they ignored them.

The lateral position of the interrupters allowed us to assess how they were processed using a suite of lateralized ERP signals. Interrupters elicited an N2pc followed by a sustained CDA when they had to be discriminated. This suggests that participants attend task-relevant interrupters, then encode them into visual working memory. Conversely, when interrupters could be ignored, there was a Pd instead of an N2pc and no CDA was observed. Thus, task irrelevant interrupters were actively suppressed from being encoded into visual WM. In contrast, spatial attention was captured regardless of whether the interrupters were task relevant, as shown by a reliable decline in alpha power contralateral to the position of the interrupters.

These findings suggest that observers could exert attentional control over whether the interrupters entered into working memory, and that this could be accomplished even when the interrupters captured spatial attention. Thus, these findings converge with prior work that has pointed towards distinct computational roles for CDA and alpha activity, with the former associated with item-based storage, and the latter associated with covert spatial attention (Hakim et al., 2019; Gunseli et al., 2019).

### Experiment 2

The relevance manipulation in Experiment 1 was a dual-task design, as it required participants to maintain information about two different tasks. With this kind of manipulation, participants could always try to optimize their performance by trading off between the two tasks on some portion of trials, especially during the relevant condition blocks. It’s possible that participants simply chose to utilize an “offline” strategy on some trials in which they did not attempt to actively maintain the target items so that they could dedicate resources to the discrimination task. This strategy may be less likely in the irrelevant condition when subjects know they do not need to do anything with the interrupting items. Such a difference in strategy could plausibly explain why we observe a CDA to the distractors for the relevant condition and not for the irrelevant condition. It is also generally consistent with our finding that the behavioral deficit was largest for the relevant condition. If this were the case, it would suggest that the results of Experiment 1 were the result of general strategic differences between the conditions that occur prior to interruption rather than the impact of the relevant interruption itself. Therefore, in Experiment 2, we tested whether participants in the relevant condition actively encoded the target items into working memory prior to the onset of the interrupters, or whether they chose not to actively encode or maintain the target array in anticipation of making the discrimination. Therefore, the key question in Experiment 2 is whether there are differences in the CDA and alpha power lateralization between the ignore and discriminate conditions during the pre-interruption period. If participants did not actively store the memory array items in WM pre-interruption, the CDA should be reduced or eliminated in the relevant condition as compared to the irrelevant condition.

## Materials & Methods

### Participants

Twenty-nine volunteers, naïve to the objective of the experiment participated for payment ($15 USD per hour). The data from 8 participants were excluded from the analysis because of too many artifacts (same criteria as Experiment 1). The remaining 21 participants (9 male) were between the ages of 30 and 18 (M = 22.1, SD = 3.9).

### Apparatus and stimuli

The apparatus was identical to Experiment 1. Stimuli (Figure 4) were also identical to Experiment 1 with the following exceptions. We were interested in the neural representations of the memory array items. Therefore, we presented the memory array items laterally and the interrupting items centrally. Thus, CDA amplitude can be interpreted as encoding and maintenance of the memory array and lateralization of alpha power can be interpreted as a shift of attention towards laterally presented memory items. Therefore, at the beginning of the experiment, a horizontal diamond comprised of a green (RGB = 74, 183, 72; 52.8 cd/m^2^) and a pink (RGB = 183, 73, 177; 31.7 cd/m^2^) triangle appeared on the vertical midline 0.65° above the fixation dot. In 50% of the trials, the pink triangle pointed to the left side and the green triangle pointed to the right side, in the remaining 50% of the trials this was inverse. Half the participants were instructed to attend the hemifield that the pink triangle pointed to, and the other half was instructed to attend the hemifield to which the green triangle pointed. Memory displayed showed an array of 3 colored squares in each hemifield. Within each hemifield, there were one or two squares in the upper quadrant and two or one square in the lower quadrant. Squares could appear within an area of the display subtending 6° to the left or right of fixation and 3.1° above and below fixation. The interruption display showed four colored squares of the same size as the ones from the memory display along the midline of the screen, drawn from the remaining colors. These interrupting items were shown on the vertical midline with a randomly jittered horizontal offset of maximally 0.55° (half of an object). Probe displays showed one colored square in each hemifield in the same location as one of the squares, randomly picked, in the original array. The color of the square in the unattended hemifield was the same as the original square on 50% of trials, and different on the other 50% of trials.

**Figure 4.**
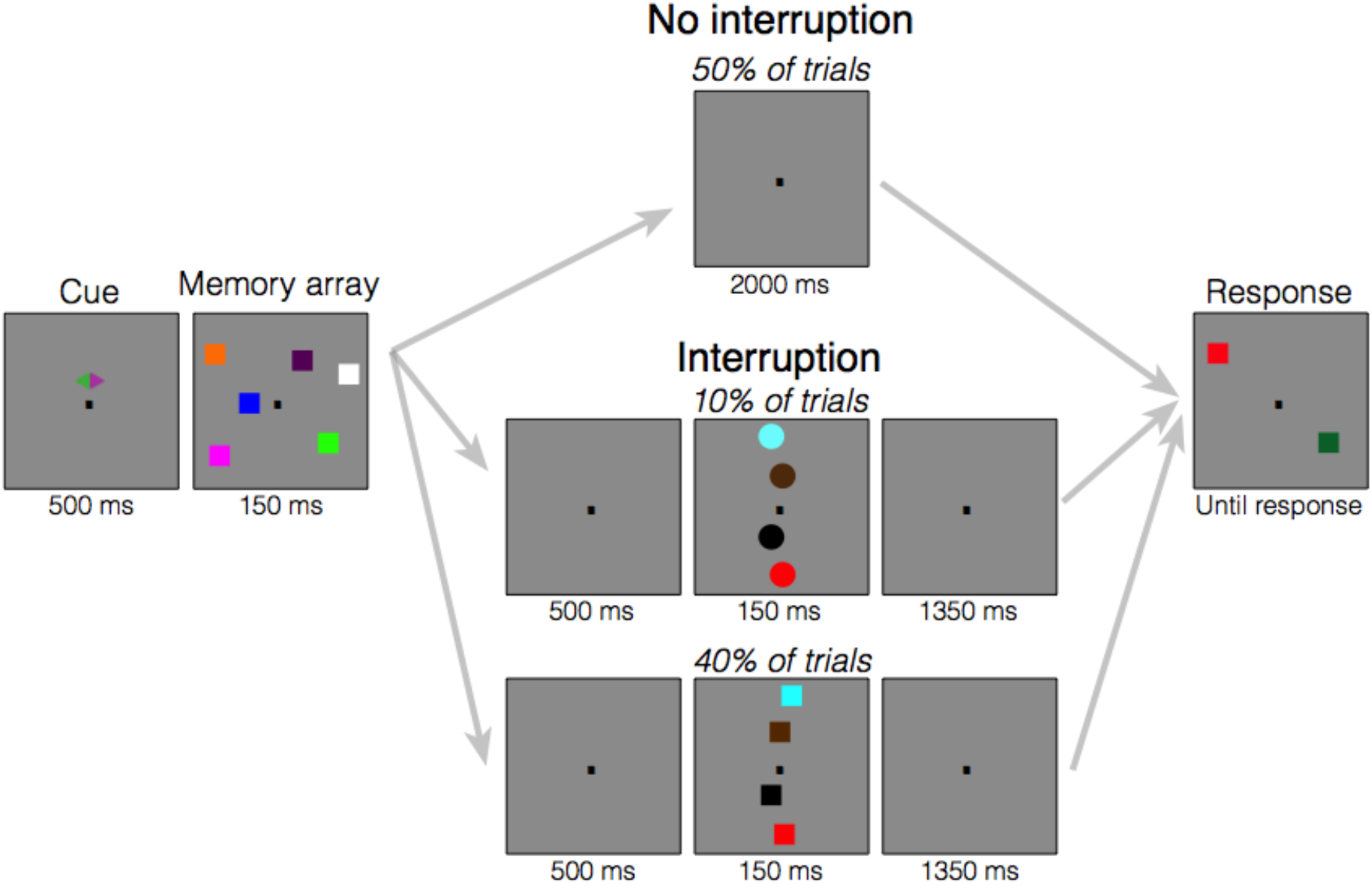
Task design for Experiment 2. At the start of each trial, a cue appeared above the fixation dot, which indicated to participants which side of the screen they should attend. Participants either attended to the green or purple side (counterbalanced across participants). Following the cue, the memory array appeared, which consisted of three colored squares on each side of the screen. Participants were told to remember the colors of the squares on the cued side. Following the memory array, the screen went blank. Then, either the screen remained blank the entire delay (no interruption conditions) or the screen went blank for 500 ms (interruption conditions) followed by a series of four objects (circles or squares) along the midline. When participants were in the “Ignore” condition, they were told to always ignore these interrupting objects. When they were in the “Discriminate” condition, they were told to determine the shape of the stimuli in order to report whether the stimuli were circles. They were told to withhold their response until the response screen appeared. Following interrupters, the screen then went blank for the rest of the delay. On the final screen, one square on either side of the screen re-appeared and could either be the same color as the original square or it could be a different color that did not appear in the display. In both conditions, participants had to report whether the square on the cued side of the screen changed colors. In the “Discriminate” condition, participants additionally had to report whether the interrupters were circles, if there were interrupters on that trial.

### Procedure

The procedure was identical to Experiment 1 with the following exception. Each trial began with a cue display (500 ms) indicating the to-be-attended side of the screen (left or right).

#### Artifact rejection & analyses

Artifact rejection and analyses were identical to Experiment 1 with the following exceptions. Since circle interrupters were only 10% of interrupted trials and required an additional response, we only included square interrupter trials in all analyses. Additionally, we analyzed EEG data in all four conditions because stimuli were presented laterally in all cases. We compared conditions with a repeated measures ANOVA with the within-subjects factors Interruption (interruption versus no interruption) and Relevance (ignore versus Discriminate). For alpha power, we analyzed all of the same time windows as the CDA because previous research has directly investigated the time course of alpha lateralization following centrally presented interruptions (Hakim et al 2020). Finally, we did not analyze the Pd and N2pc because interrupters were presented centrally in this experiment.

## Results

### Behavior

Participants remembered fewer items when they were interrupted (M=1.62±0.66) than when they were not interrupted (M=1.94±0.65), significant main effect of Interruption (F(1,20)=100.21, p<0.001, η_p_^2^ =0.83). Participants also remembered fewer items when they had to discriminate the interruptions (M=1.69±0.64) than when they could be ignored (M=1.87±0.63), significant main effect of Relevance (F(1,20)=10.20, p=0.005, η_p_^2^ =0.34). The difference between trials with and without interrupters was significantly larger when the interrupters had to be discriminated (ΔM=0.479±0.221) than when they could be ignored (ΔM=0.177±0.258), significant interaction of Interruption and Relevance (F(1,20)=13.67, p=0.001, η_p_^2^ =0.406). Additionally, the significant follow-up t-test showed that the difference between trials with and without interrupters was significantly larger in the Ignore than the Discriminate condition (t(20)=-3.698, p=0.001). Behavioral results from Experiment 2 are depicted in Figure 5.

**Figure 5.**
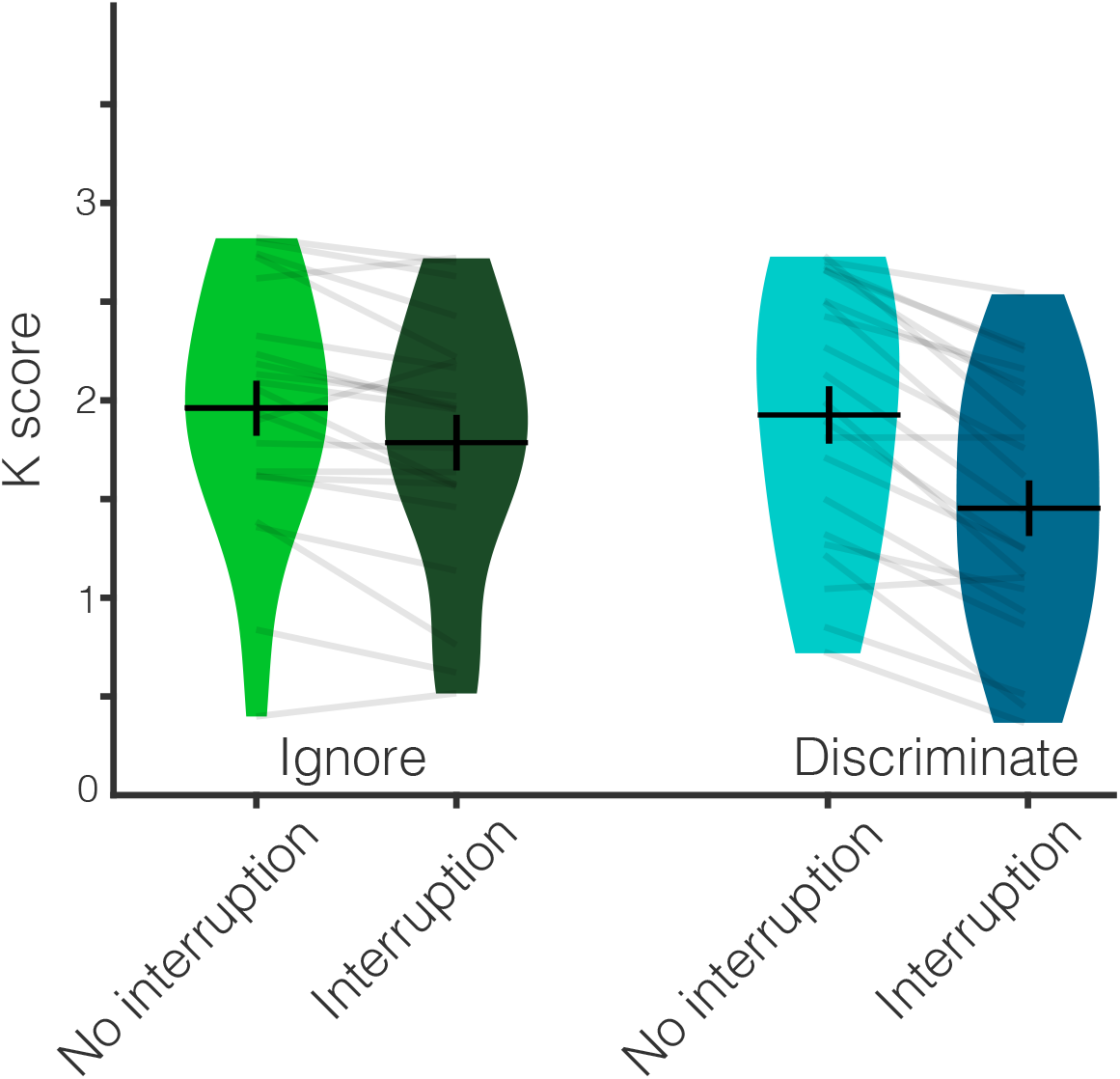
Behavioural results from Experiment 2. Behavioural performance (K score) across the four conditions. Participants remembered fewer items when they were interrupted than when they were not interrupted. This impact of interruption was larger when participants had to discriminate the interrupters than when they could ignore them. Average K score is represented by the horizontal black line and the black error bars reflect the standard error of the mean. The distribution of K scores in each condition for all participants is represented by the violin plots. Light gray lines connect data from one participant across conditions.

### Lateralized ERP

Just as in Experiment 1, we ran a repeated measures ANOVA with the factors Interruption (no interruption, interruption) and Relevance (ignore, discriminate) to determine whether there were any differences between conditions at each time point. In this experiment, we analyzed lateralized alpha power and CDA. To determine whether the signals were significantly lateralized, we additionally calculated one-way t-tests for each condition. Results displayed in Figure 6.

**Figure 6.**
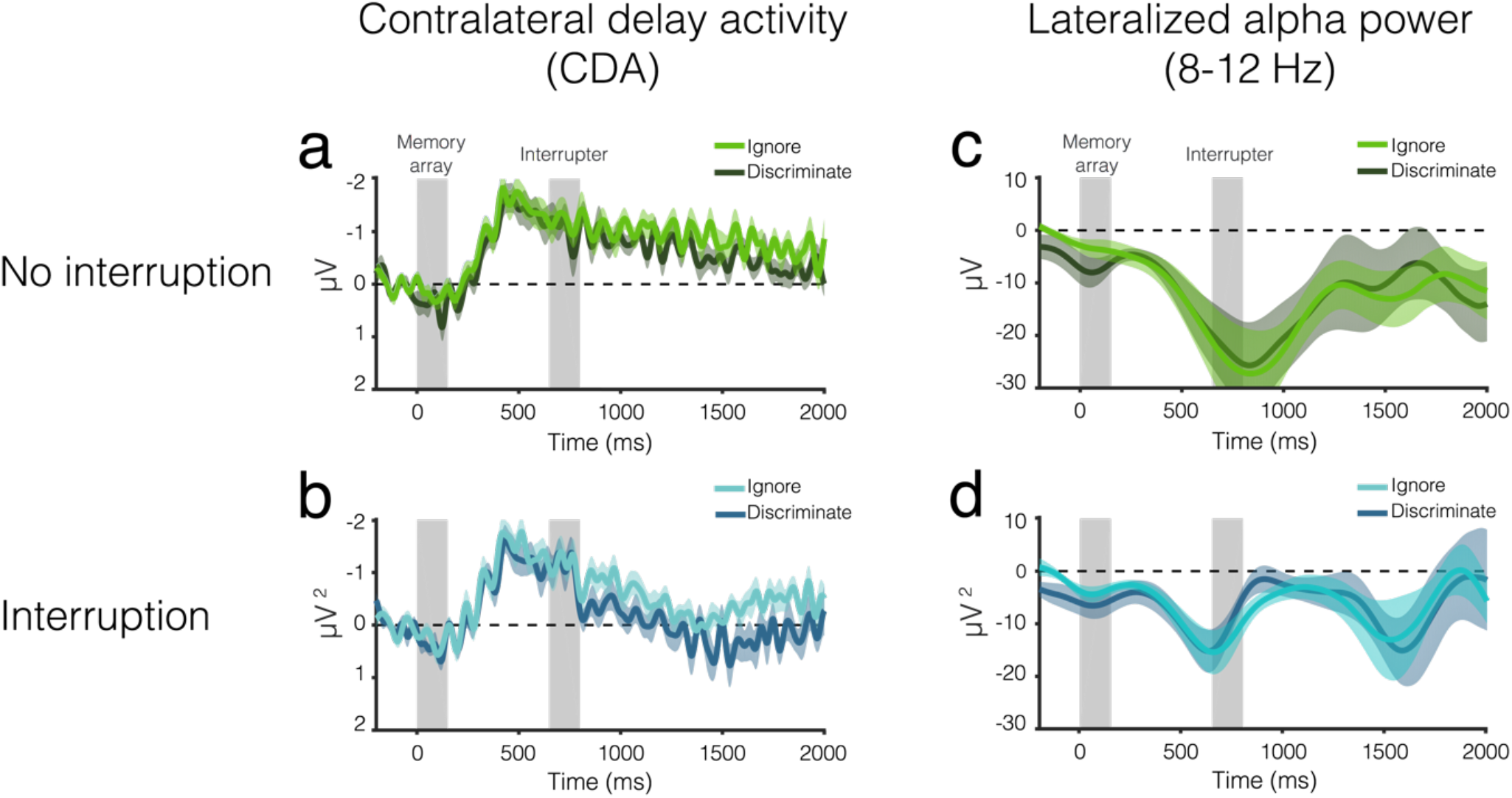
EEG results from Experiment 2. CDA amplitude over time for trials (b) with and (a) without interruption. The light color envelopes around each line represent standard error of the mean for each condition. The first vertical gray bar (0-150 ms) represents when the memory array was on the screen, and the second gray bar (650-800 ms) represents when the interrupters were on the screen, if there were interrupters on that trial. Lateralized alpha power over time for trials (d) with and (c) without interruption.

#### Pre-interruption (400-650 ms)

The CDA was reliable in all conditions (all one-sample t-tests p ≤ 0.001), and CDA amplitude did not vary between conditions (main effect of Interruption and Relevance and their interaction all p ≥ 0.24).

#### Post-interruption I (850-950 ms)

CDA amplitude was larger on trials when interrupters could be ignored (M=-0.90±0.76) than on trials when interrupters had to be discriminated (M=-0.59±0.79) regardless of whether interrupters appeared (significant main effect of Relevance: F(1,20)=2.08, p=0.02, η_p_^2^ =0.23). The main effect of Interruption and the interaction of Interruption and Relevance were not significant (both p≥0.12).

#### Post-interruption II (1050-1300 ms)

CDA amplitude was larger on trials without interrupters (M=-0.83±0.51) than on trials with interrupters (M = −0.22 ± 0.73) regardless of relevance (significant main effect of Interruption: F(1,20) = 17.73, p < 0.001, η_p_^2^ = 0.47). The main effect of Relevance was trending, but not significant (F(1,20) = 3.65, p=0.07, η_p_^2^=0.15), and the interaction of Interruption and Relevance was not significant (F(1,20)=0.29, p = 0.69, η_p_^2^ = 0.01). When interrupters were not present, there was a reliable CDA regardless of condition (both p ≤ 0.001).

### Lateralized alpha power

#### Pre-interruption (400-650 ms)

Before the interrupters appeared (400-650 ms), alpha power was significantly lateralized in all conditions (all p≤0.004), and lateralization did not vary between condition (main of Interruption and Relevance and their interaction were not significant all p ≥ 0.25).

#### Post-interruption I (850-950 ms)

Immediately following interruption (850-950 ms), alpha power was more lateralized on trials without interruption (M=-25.39±33.33) than on trials with interrupters (M=-3.71±6.89) regardless of condition (significant main effect of interruption: F(1,20)=9.37, p=0.006, η_p_^2^=0.32). The main effect of Relevance and the interaction of Relevance and Interruption were not significant (both p ≥ 0.42).

#### Post-interruption II (1050-1300 ms)

Toward the end of the delay (1050-1300 ms), alpha power was more lateralized on trials without interrupters (M=-13.66±24.62) than on trials with interrupters (M=-3.77±12.11; significant main effect of interruption: F(1,20)=5.98, p=0.02, η_p_^2^=0.23). The main effect of Relevance and the interaction of Relevance and Interruption were not significant (both p ≥ 0.98). Follow-up one-sample t-tests revealed that alpha power was significantly lateralized on trials without interruptions (both p ≤ 0.03), but not on trials with interruptions (both p≥0.09), regardless of relevance. To summarize, the presentation of the central distractors disrupted alpha lateralization towards the target locations, and this effect did not depend on whether the distractors were task relevant.

### Conclusions

In Experiment 2, we were interested in whether interrupter relevance affected the likelihood of encoding and maintaining target information in WM. The relevant interruption condition in our two experiments is similar to a dual-task design as participants have to maintain information about two separate tasks. Participants could decide to drop information about the initial target array in order to encode the new information from the second task. Alternatively, they could attempt to sustain the initial working memory representations at the expense of sufficiently attending the relevant interrupters. How did participants address this trade-off between tasks in the present study?

Before the interrupters appeared, there was no difference in CDA amplitude or alpha power lateralization between the ignore and discriminate conditions. There was a clear CDA and alpha power lateralization for both conditions, with no differences between them. This indicates that in both conditions, participants encoded and maintained lateralized working memory representations and sustained their attention. Differences between conditions only emerged after the time when interrupters were supposed to appear. Therefore, the results from Experiment 1 are not simply due to a dual-task tradeoff between the encoding and maintenance of the memory array and the interrupters.

The interrupters always appeared at the same time during the delay. Therefore, participants could anticipate when they might be interrupted. Around the time when interrupters typically appeared, the CDA was smaller when participants anticipated a relevant interrupter than when they anticipated an irrelevant interrupter. During this same time window, alpha power lateralization depended on the presence of interrupters. Towards the end of the trial, however, both CDA amplitude and alpha power lateralization were closer to baseline when interrupters were present than when they were not, regardless of relevance. Overall, when participants anticipate that they may have to integrate new information into working memory, they hold less information about the target array around the time that they think new information will appear, even if this new information doesn’t appear. However, by the end of the trial, relevance no longer impacts the likelihood of sustaining attention or maintaining information about the target array. Overall, we can rule out the alternate explanation of experiment 1 that participants prematurely drop information about the target array when relevant interrupters are presented. Participants were clearly attending and maintaining memoranda even when relevant interrupters appeared.

## Discussion

The key finding of the present study was that processing of stimuli that interrupt ongoing WM representations depended on their relevance. Relevant interruptions were encoded and maintained in WM, whereas irrelevant interruptions were suppressed and never entered WM. On the other hand, spatial attention was captured regardless of stimulus relevance. In Experiment 2, we investigated whether participants in the relevant condition of Experiment 1 actively encoded and maintained memory items prior to the onset of the interrupters. We found that pre-interruption, there were no differences between conditions. Participants encoded and maintained working memory representations and sustained attention to the memory items equally in both conditions. Differences between conditions only emerged after interruption onset, indicating that the results from Experiment 1 are not solely driven by a dual-task tradeoff between the maintenance of the memory array and the interrupters.

We observed distinct effects of interrupters on lateralized ERP signals and lateralized alpha power in these experiments. The results converge with past proposals of a distinction between item-based and spatial capture of attention (Hakim, Adam et al. 2019). In our procedure, alpha oscillations showed that interrupters captured spatial attention, regardless of task relevance. By contrast, the formation of item-based representations in working memory was completely determined by task relevance, such that encoding into working memory was suppressed when observers could ignore the interrupters. Interestingly, both item-based storage and spatial attention towards the memoranda were eventually disrupted by the presentation of the interrupters, regardless of whether a dual task was imposed.

### Spatial capture by relevant and irrelevant interrupters

In Experiment 1, we directly investigated how interrupters are processed. In this experiment, we found that alpha power was significantly lateralized following the onset of both relevant and irrelevant interrupters. This result provides evidence that spatial attention shifted to the location of both the relevant and irrelevant interrupters. We designed our interrupters to be very similar to the original memoranda in order to induce a large behavioral deficit on interrupted trials compared to non-interrupted trials. Previous research has shown that when interrupters share features with memoranda (Hollingworth & Beck, 2016; Olivers & Eimer, 2011; Soto et al., 2008; van Moorselaar et al., 2015) or are part of the attentional set (Charles L. Folk et al., 2008; Charles L. Folk & Remington, 1998), spatial attention can be captured at the location of the interrupters. However, the similarity between targets and interrupters is a continuum (Duncan & Humphreys, 1989) and attentional capture increases with an interrupter’s similarity to a target (Ansorge & Heumann, 2003; Ludwig & Gilchrist, 2002). In the present study, relevant interrupters were the same shape as the targets and drew their color from the same pool of colors as the targets, i.e., they had potential target colors. However, no interrupter was ever the same color as a target within a trial. Nevertheless, the finding that spatial attention was captured by both types of interrupters may be partially due to the fact that even irrelevant interrupters were perceptually similar enough to the targets (i.e. contingent capture). Future research should determine whether spatial attention is necessarily captured when interrupters do not share any similarities with currently maintained working memory representations. Understanding whether spatial and item-based attention are captured along the entire continuum is an important question as it will give insight into the question of how features are weighted in WM and how this affects attention deployment.

### Voluntary control of item-based capture

In Experiment 1, irrelevant interrupters elicited a Pd, suggesting that they were actively suppressed. Relevant interrupters, however, first elicited an N2pc which indicates that these items were individuated. The subsequent lateralized negativity then transitioned into a CDA, suggesting that distractors were not only individuated but also encoded into WM. These results illustrate that participants dynamically respond to task demands by suppressing irrelevant interrupters from WM and only encoding relevant interrupters into working memory. Even when salient, bottom-up stimuli captured spatial attention, participants still had voluntary control over whether to store that information in working memory. These results are in line with previous findings showing that successful suppression of irrelevant information can contribute to better performance (Feldmann-Wüstefeld et al., 2016; Gaspar & McDonald, 2014a; Risa Sawaki & Luck, 2012; Weaver et al., 2017). For example, when target identity is correctly reported in a visual search task, a concurrently presented salient distractor elicits a pronounced Pd component, indicative of active suppression (Feldmann-Wüstefeld et al., 2020). Conversely, when the distractor identity is erroneously reported, distractors elicit a CDA and a less pronounced Pd, suggesting that the distractor was encoded into WM.

Our results also nicely align with the working memory gating literature. This literature provides a framework to explain which information is allowed to enter working memory and which information is blocked (Badre, 2012; Chatham et al., 2014; Chatham & Badre, 2015; O’Reilly & Frank, 2006). According to this account, the working memory gate is the mechanism by which irrelevant information is blocked from entering. When the working memory gate is open, it allows information to enter working memory. When it is closed, ongoing working memory representations are sustained, while irrelevant information is blocked (Badre, 2012). The working memory gating literature has mostly used fMRI to demonstrate which parts of the brain are involved in working memory gating and maintenance and has not distinguished between WM gating and capture of spatial attention. Our results suggest that we may be able use EEG activity to track working memory gating as well. We propose that the CDA tracks how much information passes through the gate, whereas the Pd reflects the gate itself. Previous research has shown that Pd amplitude scales with the number of items that were blocked from entering working memory (Feldmann-Wüstefeld et al., 2019). Therefore, Pd amplitude may reflect how firmly the working memory gate was closed. Conversely, the CDA could reflect how much information is encoded into working memory, with relevant information more likely to pass through the gate. Based on our results, top-down control may determine how firmly we close the gate, whereas stimulus salience or relevance determine how much information enters working memory. Future research could investigate the precise temporal dynamics of working memory input and output gating using these proposed EEG signals. For example, if the working memory gate accidentally allows irrelevant information into working memory, how long is that information maintained in working memory before it is dropped?

### Is attentional capture obligatory?

Our findings provide a new framework in which we can investigate attentional capture. We propose that attentional capture is comprised of item-based and spatial capture. Item-based capture involves forming working memory representations of the interrupting stimuli. Based on our findings, item-based capture appears to be subject to voluntary attentional control. It allows relevant stimuli to enter working memory, while irrelevant stimuli are suppressed. We also have clear evidence that, in our specific task context, spatial capture occurred when interrupters were present, regardless of top-down goals. Thus, WM gating can successfully block irrelevant information, even when spatial attention is captured. From our perspective, information needs to be held in working memory in order for it to be processed. Therefore, if interrupting stimuli are not encoded into working memory via item-based capture, they are not fully processed, even if they captured spatial attention. Selecting an item using spatial attention is necessary, but not sufficient, to encode it into working memory.

We investigated item-based and spatial capture during maintenance of ongoing working memory representations. However, the majority of the attentional capture literature has investigated this process during encoding. Therefore, future research could apply this new framework of attentional capture to investigate item-based and spatial capture during encoding. This could potentially provide insight into the ongoing debate about whether attentional capture during encoding is obligatory (Feldmann-Wüstefeld et al., 2020; Feldmann-Wüstefeld & Schubö, 2013; Gaspar & McDonald, 2014b; Gaspelin et al., 2015, 2016; Gaspelin & Luck, 2018; Hickey et al., 2009; Liesefeld et al., 2017; R. Sawaki et al., 2012). We hypothesize that the ways in which the two sub-components of attentional capture (item-based and spatial capture) respond during encoding should be similar to how they respond during maintenance. That is, we hypothesize that during encoding, spatial capture may happen regardless of top-down goals, whereas item-based capture may be subject to voluntary attentional control.

### Impact of interrupters on ongoing working memory representations

Previous research has demonstrated that salient interrupting stimuli interfere with object representations in WM and cause attention to shift away from maintained representations (Hakim, Feldmann-Wüstefeld, et al., 2019). Our work replicates and extends these findings by adding a top-down perspective. Participants had to either attend (relevant) or ignore (irrelevant) interrupters. Here, we show that the CDA, a neural measure of working memory load, was initially influenced by top-down goals. When participants anticipated that they may have to encode additional information into WM, the CDA was smaller than when participants knew that interrupting stimuli could be ignored. However, the CDA was at baseline by the end of trials that contained both relevant and irrelevant interrupters. This suggests that both types of interrupters harmed lateralized object representations of the memoranda. However, towards the end of the trial, alpha power, a neural measure of spatial attention, shifted to baseline following both relevant and irrelevant interrupters, suggesting that participants shifted their attention away from the locations of the original memoranda regardless of whether the interrupting information was relevant. These results illustrate that the observers’ goals determine the encoding of item-based representations, even when spatial attention is captured. By the end of the trial, however, interrupters harm both spatial attention and the object representations of the memoranda, regardless of top-down goals.

### Conclusions

Previous research has treated attentional capture as a monolithic process. Here, we present new evidence that there are at least two sub-component processes of attentional capture that are neurally dissociable: spatial capture and item-based capture. Lateralized alpha power indexes spatial capture, a process that involves a shift of spatial attention. By contrast, item-based capture is tracked by the N2pc and CDA when item-based representations are deemed relevant and allowed to enter working memory, while the Pd tracks the active suppression of items from WM. This fractionation of attentional capture into distinct sub-component processes provides a framework by which the fate of ongoing WM processes after interruptions can be explained. We show that relevant interrupters trigger both of these dissociable processes which means that they undermine WM maintenance more. Irrelevant interrupters, however, only trigger spatial capture from which ongoing WM representations can recover more easily.

## Data availability

Datasets for all experiments will become available online on Open Science Framework upon acceptance of the manuscript or reviewer request.

## Acknowledgements

Research was supported by NIMH grant ROIMH087214 and Office of Naval Research grant N00014-12-1-0972.

## Author contributions

NH, TFW, and EKV conceived of the study. NH and TFW performed analyses. NH collected the data and wrote the initial draft of the manuscript, which all authors read and edited.

